# Changepoint detection versus reinforcement learning: Separable neural substrates approximate different forms of Bayesian inference

**DOI:** 10.1101/591818

**Authors:** Wolfgang M. Pauli, Matt Jones

## Abstract

Adaptive behavior in even the simplest decision-making tasks requires predicting future events in an environment that is generally nonstationary. As an inductive problem, this prediction requires a commitment to the statistical process underlying environmental change. This challenge can be formalized in a Bayesian framework as a question of choosing a generative model for the task dynamics. Previous learning models assume, implicitly or explicitly, that nonstationarity follows either a continuous diffusion process or a discrete changepoint process. Each approach is slow to adapt when its assumptions are violated. A new mixture of Bayesian experts framework proposes separable brain systems approximating inference under different assumptions regarding the statistical structure of the environment. This model explains data from a laboratory foraging task, in which rats experienced a change in reward contingencies after pharmacological disruption of dorsolateral (DLS) or dorsomedial striatum (DMS). The data and model suggest DLS learns under a diffusion prior whereas DMS learns under a changepoint prior. The combination of these two systems offers a new explanation for how the brain handles inference in an uncertain environment.

**One Sentence Summary:** Adaptive foraging behavior can be explained by separable brain systems approximating Bayesian inference under different assumptions about dynamics of the environment.

## Main Text

Everyday life constantly forces humans and other animals to predict future states of our environment, often based on limited data. For example, animals foraging for food must choose among different locations in order to achieve the highest ratio of energetic gain to cost. This decision entails a form of induction, using past experience to infer the probability of finding food at each possible location.

Bayesian inference provides a normative framework for predicting future events from past experience. Most applications of Bayesian inference test hypotheses about latent variables in the environment, which are probabilistically related to observable outcomes. In the case of foraging, this process can provide a posterior probability that food will be available at each location, given the currently available data from past visits. Bayesian models of cognition have been successful in recent years at explaining human reasoning and decision-making as optimal inference in a variety of complex cognitive tasks, including language processing and acquisition (1), word learning (2), concept learning (3), causal inference (4), and deductive reasoning (5).

However, Bayesian inference by itself is not a complete solution to inductive problems. The general problem of induction is that it is logically impossible to make predictions without committing to some a priori, experience-independent assumptions about how the world works (6, 7). For any inductive algorithm, there exist environments in which it will fail catastrophically (8, 9). Therefore, making approximately correct assumptions about which processes generate the observable data is critical for successful prediction. In the context of Bayesian inference, the classical problem of induction is formalized as a problem of selecting the right hypothesis space and prior distribution, or equivalently a *generative model*. A generative model is a specification of latent (unobservable) random variables together with assumptions about their probabilistic causal relationships to the observable data. Most Bayesian models of cognition impute rich generative models to the minds of subjects without addressing the hard question of how the brain selects them over other alternatives (10, 11).

Recent research offers two proposals for how the brain deals with uncertainty about the latent causal structure in the environment. One approach argues that Bayesian optimality is unattainable outside of “small worlds” where the space of possible generating processes is constrained (12), and that instead the brain relies on simple algorithms that are robust to this type of uncertainty (13). A second approach uses hierarchical inference over many layers of representations, with extremely abstract processes at the top layers that can generate a wide variety of different structures at the lower layers (14, 15).

We propose a third possibility, in the tradition of multiple-systems or mixture-of-experts models of learning (16, 17), in which different brain systems approximate Bayesian inference with respect to different generative models. A computational advantage of this *mixture of Bayesian experts (MBE)* architecture, relative to the second possibility just mentioned, is that the assumptions made by each system might be fairly simple and amenable to efficient approximate implementation. Under this division of labor, the brain could effectively hedge its bets and succeed in tasks that obey different types of dynamics.

We apply the MBE framework to the foraging domain, where a primary challenge for making effective predictions comes from nonstationarity of resource availability at each location. We consider two simple assumptions the brain might make about the temporal structure of this nonstationarity, both of which are suggested by previous models of learning and by neurophysiological data. First, reward probability at each location could be generated by a *diffusion process*, characterized by continuous stochastic change over time, following a variant of Brownian motion (Figure 1A). Second, reward probability at each location could be generated by a *changepoint process*, characterized by a step function with periods of stationarity interrupted by abrupt jumps (Figure 1B). Bayesian inference under these two assumptions produces qualitatively different patterns of behavior, with each outperforming the other when its assumptions are met (Figure 1C).

**Figure 1:**
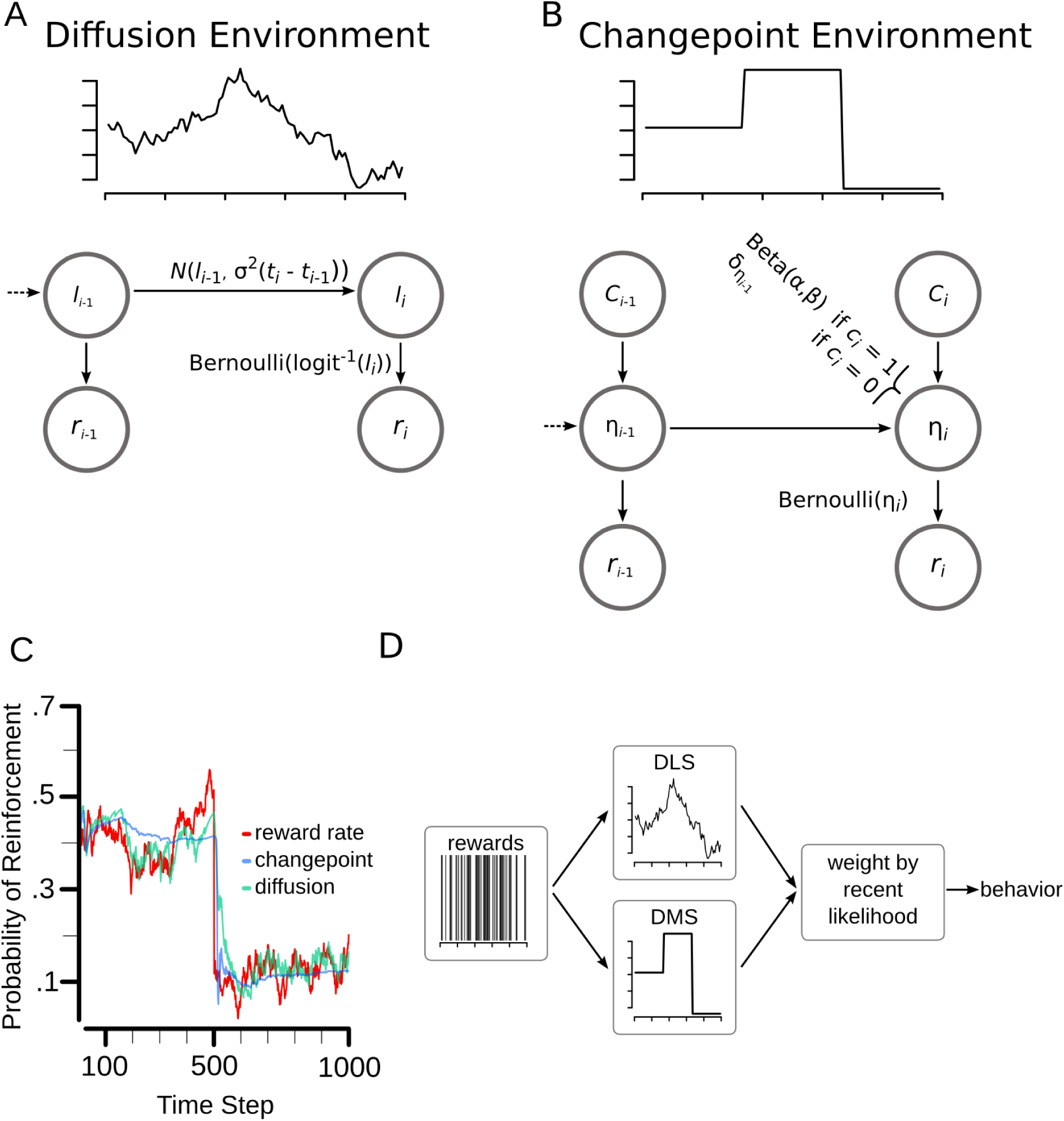
Mixture of Bayesian experts (MBE) framework as applied to a nonstationary binary reward task (see Supplementary Material for details). A. Sample trajectory of reinforcement rates in a diffusion environment, and causal graphical model of diffusion process (*l*: reward log-odds, *r*: reward, *i*: time index, σ: diffusion rate, *N*: normal distribution). B. Sample trajectory of reinforcement rates in a changepoint environment, and causal graphical model of changepoint process (*c*: changepoint indicator variable, η: reward probability, δ: Dirac delta). C. Optimal inference based on diffusion and changepoint models with simulated data from a jump diffusion process. Binary reward values were sampled on each time step from a Bernoulli distribution determined by the reward rate at that time. Curves for the changepoint and diffusion models show the posterior estimate of the reward rate at each time step, conditioned on the reward history prior to that step. The changepoint model adapts more rapidly to the changepoint, whereas the diffusion model better tracks the more continuous changes in reinforcement rate. D. Schematic illustration of full model, which combines predictions from both subsystems.

Both diffusion and changepoint processes are arguably common in foraging environments and other natural learning tasks. Whether food is available at a location may be affected by random visits from many other foraging individuals, or other small perturbations, producing diffusion dynamics. On the other hand, climate changes or other large discrete events may cause sudden changes in the availability of food, producing changepoint dynamics. Economic models of prices in financial markets have also converged on these two classes of stochastic processes as important for modeling nonstationarity (18).

Two broad classes of models for human and animal learning correspond closely to inference under the assumptions of diffusion and changepoint environments. First, models founded on correcting prediction error, including classical models of human probability learning (19) and animal conditioning (20), as well as the modern computational framework of reinforcement learning (21), implicitly assume continuous random drift in environment parameters and thus approximate optimal inference with respect to a diffusion process. Second, recently developed models of sequential effects in learning (22, 23) and of learning in artificial changepoint tasks (24, 25) are explicitly cast as Bayesian inference assuming a changepoint process. Algorithmically, these models can be implemented by particle filters that maintain hypotheses about the current state of the environment, with hypotheses (particles) being abandoned when they are too inconsistent with new observations (24). Thus these models are also similar to older hypothesis-testing models such as win-stay lose-shift (26).

Existing behavioral neuroscience data suggest the brain may have developed separable systems supporting these two approaches to learning in nonstationary environments, each involving a distinct subregion of dorsal striatum (27–29). The first of these regions, dorsolateral striatum (DLS), is interconnected with motor cortices and is thought to gradually learn the probability of reward associated with different response alternatives through reinforcement learning (30, 40). The second region, dorsomedial striatum (DMS), is interconnected with prefrontal cortical (PFC) areas involved in working memory. Computational models of the interaction of these two regions suggest DMS gates the updating of working memory representations in PFC (31), thus implementing the type of discrete hypothesis updating needed for changepoint detection (24). We therefore hypothesized that the dorsal striatum participates in an implementation of the MBE architecture proposed above, with the DLS approximating Bayesian inference under the assumption of diffusion dynamics and the DMS approximating Bayesian inference assuming changepoint dynamics.

We formalized this proposal as an idealized computational model comprising separate systems performing these two forms of Bayesian inference (see Supplementary Material). Each system maintains estimates of the current reward probability for every available action, and action selection is guided by these estimates. We predicted that when DMS functioning is impaired, behavior will be well described by the diffusion model alone (i.e., by inference assuming a diffusion process in the environment), and that when DLS functioning is impaired, behavior will be well described by the changepoint model alone. In an intact subject, we predicted both systems to contribute to behavior. In this case we assume normative integration of the predictions from the two subsystems, which entails a hierarchical model in which each subsystem’s prediction is weighted according to how well it has predicted recent outcomes (formally, likelihood-weighted averaging with a recency bias; see Figure 1D and Supplementary Material). This integration would allow the brain to infer the relative prevalence of each type of dynamics in a given context and to act accordingly.

## Results

We applied our framework to data from a recently published study indicating that DLS and DMS play distinct roles in adaptation to changes in reward contingency (27). In this experiment, rats were trained to press two levers for reward on a concurrent variable-interval (VI) schedule, in a laboratory analog of foraging environments. The VI scheduled was defined such that, after each time a reward was received from pressing a lever, that lever was nonrewarding for a variable amount of time, drawn from a uniform distribution ranging from 0 to *T* seconds. In phase 1 of the experiment, rats were trained on the same set of contingencies for 6 consecutive sessions. During this training, one lever followed a more favorable schedule than the other (*T* = 20 s vs. 80 s). One day after the last training session, either the zeta inhibitory peptide (ZIP) or saline was infused into either the DMS or the DLS. Previous research suggests that ZIP infusion should erase any task-relevant memory from the infused region, by inhibiting the putative memory-maintenance protein kinase C isozyme protein kinase M**ζ** (PKM**ζ**) (32).

Two days after the infusion, the rats were tested on their adaptation to a contingency change (phase 2). The lever that had been associated with the more favorable schedule was now associated with the less favorable schedule, and vice versa. The results of the experiment showed that if ZIP was infused into the DMS, rats adapted to the contingency shift more slowly than did the saline controls. On the other hand, if ZIP was infused into the DLS, rats adapted to the contingency shift more rapidly than did the controls (Figure 2A).

**Figure 2:**
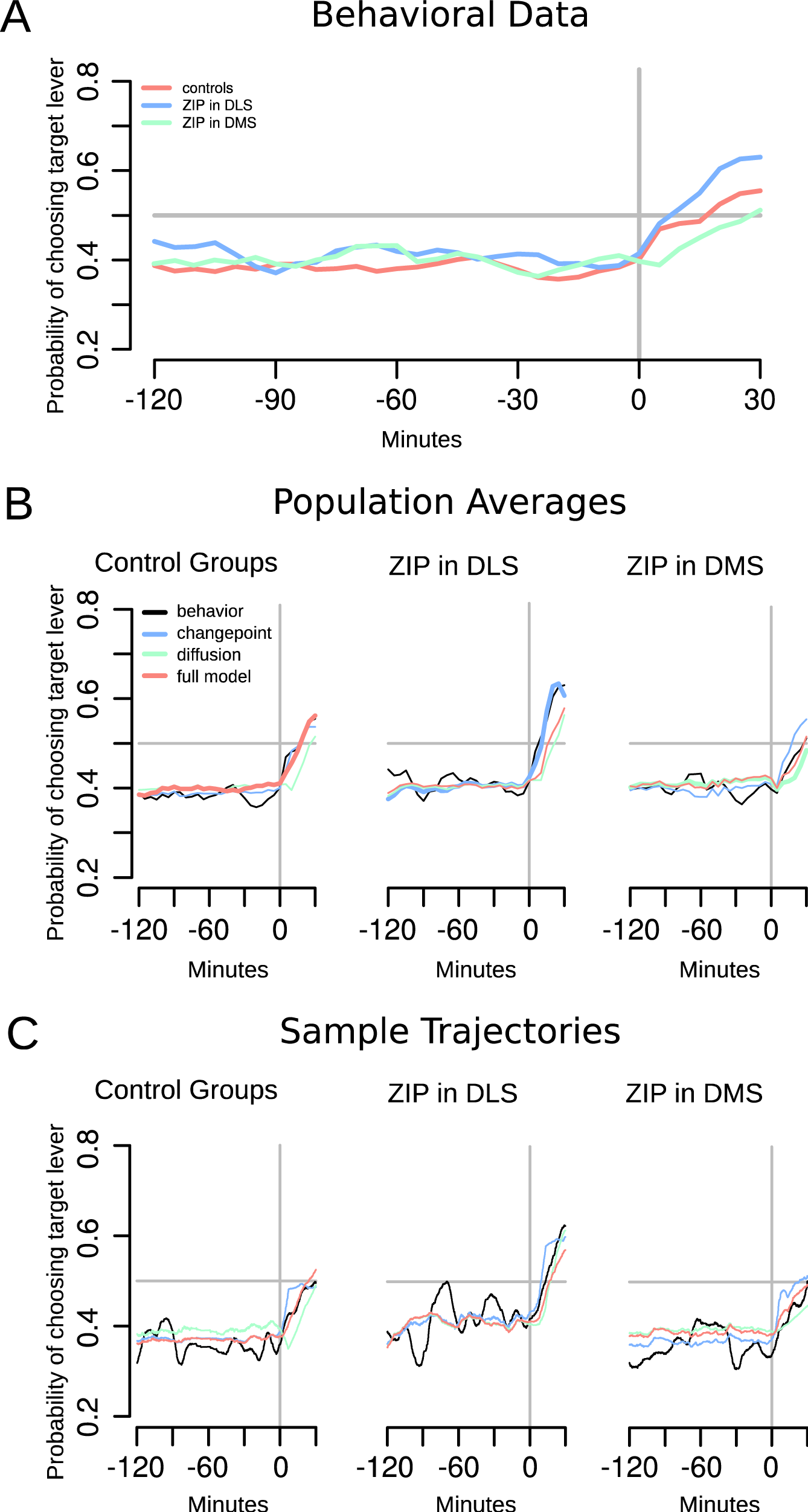
Behavioral data from Ref. 27 and model predictions. A. Population averages of behavioral data (Gaussian filtered). Vertical line at time 0 separates phases of the experiment, when infusions were administered and the target lever switched from being less to more rewarding. Animals that received ZIP infusions in the DLS adapted more rapidly than animals with control infusions, and animals with DMS infusions adapted more slowly. B. Population averages of behavioral data (same as in A) and averaged model predictions for the three groups. DMS group is best fit by the diffusion model, DLS group by the changepoint model, and control groups by the full model. C. Same as B but for a single animal from each group.

We hypothesized that these differences in the rats’ rates of adaptation are the result of the ZIP-DMS group behaving more in accordance with diffusion-based inference, and the ZIP-DLS group behaving more in accordance with changepoint-based inference (cf. Figure 1C). Specifically, microinfusions of ZIP into either dorsal striatal region should cause that region to “forget” how to support reward prediction in this task (27). According to the model, the poor predictive performance of this system causes its predictions to be effectively ignored due to the likelihood-weighted averaging in the MBE framework. Thus behavior is dominated by the predictions of the other (intact) system.

To test this explanation, we fit each of three models to the complete sequence of lever presses of each individual rat from phases 1 and 2 (see Supplementary Material for details). The full model assumed that both diffusion-and changepoint-based inference systems were operational throughout the experiment, with their predictions combined by likelihood-weighted averaging to determine behavior. The diffusion model assumed both systems were intact and influencing behavior in phase 1, but that behavior in phase 2 was controlled by the diffusion system alone (to simulate the effects of microinfusion to the DMS). Likewise, the changepoint model assumed both systems were intact in phase 1, but that behavior in phase 2 was controlled by the changepoint system alone (to simulate the effects of microinfusion to the DLS). Although the models differed only in phase 2, it was important to simulate them on both phases so that the prior distributions at the start of phase 2 were determined by the animal’s experience during phase 1.

As predicted, the ZIP-DLS group was fit best by the changepoint model, the ZIP-DMS group was fit best by the diffusion model, and the control groups were fit best by the full model (Table 1 & Figure 2B). Data and model predictions for example subjects in each group are shown in Figure 2C.

**Table 1:** AIC values of model fits, summed over subjects in each group. The ZIPDLS group was fit best by the changepoint model, the ZIPDMS group was fit best by the diffusion model, and the control groups were fit best by the full model.

| Model | Group |  |  |
| --- | --- | --- | --- |
|  | ZIP-DMS | ZIP-DLS | Control |
|  | **17757.** | 22365.3 | 39437.8 |
| Diffusion | **82** | 8 | 8 |
|  | 17788.8 | **22315.** | 39511.2 |
| Changepoint | 7 | **57** | 2 |
|  | 17782.9 | 22458.2 | **39328.** |
| Full | 7 | 3 | **86** |

## Conclusions

Any computational process for learning, or for predicting future outcomes from past experience, will only be successful insofar as it matches the statistical structure of the environment in which it is implemented. Bayesian formulations of learning make this dependence explicit, but it is just as true for models specified directly at an algorithmic level. The question of how the brain develops or selects hypotheses about the structure of the environment has received much less attention than question of how learning progresses given those hypotheses. Here we have proposed the brain follows a divide-and-conquer strategy, as formalized in the MBE framework, in which different brain systems approximate optimal inference with respect to different generative models. The assumptions of each system might be quite simple, as with the diffusion and changepoint dynamics for action-reward contingencies in the foraging model described here.

It is interesting to note that, at a fine temporal scale, neither component of the present model matches the actual structure of the task environment in the experiment modeled here. The uniform distribution used in the VI schedule causes the reward probability for each action to increase linearly as a function of time since the last reward, a dependence not captured by either the diffusion or the changepoint submodel. Previous analysis of behavior in concurrent VI tasks indicates that animals are not sensitive to this type of local timing information (33), thus supporting the assumptions of our model. These observations highlight the fact that probabilistic rational models of human or animal behavior need not be founded on the actual statistical structure of a task environment, but might be more fruitfully applied to what the brain (or a subsystem) believes that structure to be.

The hierarchical portion of the present model may also explain previous data showing that rats adapt to experimenter-induced changepoints more quickly if past changepoints were encountered frequently than if reward contingencies were constant for several training sessions (25). According to the model, past changepoints increase the predictive performance of the changepoint-inference system relative to the diffusion-inference system, so that more weight is given to the former system in subsequent decision-making.

Part of the value of this framework lies in the linkage of computational, algorithmic, and implementational explanations (34). Specifically, it offers a computational-level perspective on the differential roles of different brain systems, in that they carry out algorithms that approximate distinct optimization tasks defined by alternative environmental assumptions. This perspective may also offer insight into other multiple-systems theories that postulate separation between an explicit, rule-based system and a more flexible implicit system (16). The explicit system can be cast as embodying an expectation of rule-like structure in the world, in that meaningful stimulus distinctions are aligned with perceptual dimensions and that categories or natural kinds are defined by conjunctions or disjunctions of criteria on these dimensions. The implicit system expects the world to be carved up more arbitrarily, which can explain why that system is more flexible but generally learns more slowly (35).

The present view can be contrasted with other theories in which different brain systems have identical computational goals but differ in their algorithmic approximations to those goals. Daw et al. (36) propose one such theory, cast in a reinforcement learning framework, in which both systems estimate values of actions assuming the environment is a Markov decision process (37), but one system derives estimates using temporal-difference learning and the other uses model-based lookahead. The difference between computational-and algorithmic-level competition between systems also has implications for optimal integration of their predictions. In Daw et al.’s model, the two systems make predictions under the same assumptions but using different approximations, and thus the predictions are optimally combined according to which approximation is more certain (precision-weighted averaging). In the present model, the systems make predictions under different assumptions (diffusion vs. changepoint dynamics), and thus they are weighted according to which assumption is better supported by recent data (likelihood-weighted averaging).

We have purposely cast the modeling here at a computational level, to highlight how differences in computational goals (i.e., optimal inference with respect to different generative models) can explain differential functioning of different brain systems. This does not imply an assumption that the brain performs exact inference. An important step for further work will be to implement and test algorithmic approximations of this idealized model. The diffusion subsystem should be well approximated by a reinforcement learning model that maintains a value estimate for each action (rather than a full posterior distribution) and updates that estimate toward the outcome observed each time that action is chosen (38). The changepoint system might be approximated by a PFC-gating model (31, 39), in which the PFC holds multiple hypotheses about the current reward probability for each action, and the DMS maintains or abandons hypotheses depending on their agreement with observed outcomes. A complete understanding of how the brain learns in nonstationary environments must include such algorithmic explanations. Nevertheless, the computational-level analysis offered here provides a powerful means for studying the functional organization of neural systems for learning.

## Acknowledgements

This research was supported by Air Force Office of Scientific Research grant FA9550-10-1-0177. Thanks for helpful discussions of this work go to the members of the computational cognitive neuroscience lab at the university of Colorado at Boulder, and members of the decision neuroscience lab at the California Institute of Technology.

## Supplementary Methods

The models all assume the rat chooses which lever to press based on estimates of each lever’s current probability of producing a reward. These probabilities are estimated separately for each lever, by Bayesian inference from the outcomes of past presses of that lever.

Let (*t*_*i*_) be the sequence of times *t* at which the rat presses either lever, *r*_*i*_ be the lever pressed at time *t*_*i*_ (*L* for left or *R* for right), and *x*_*i*_ an indicator of whether a reward was received (1 or 0). When selecting a response at time *t*_*n*_, the value of each lever is estimated as that lever’s current reward probability, conditioned on the history of the task:

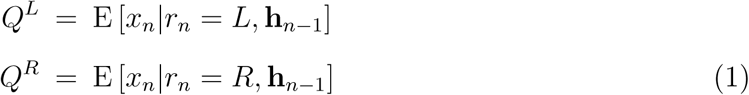

where **h**_*n-*1_ = (*r*_*i*_, *x*_*i*_) *i<n* is the history prior to *t*_*n*_.

The action is then selected according to a standard softmax rule (Luce, 1959; Sutton & Barto, 1998):

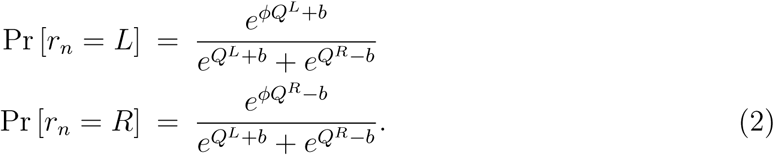

The inverse-temperature parameter, *ϕ*, controls the degree of exploration or stochasticity in behavior. The bias parameter, *b*, captures variability in individual subjects’ enduring preferences between levers.

The models thus predict which lever the rat will press (*r*_*n*_), conditioned on when the rat makes a response (*t*_*n*_). Additional models were evaluated that predict response timing in addition to response selection, by incorporating a null response (*N)* in Equation 2 with constant value *Q*^*N*^ = *u* (a free parameter), and evaluating model predictions at all time points (i.e., every 1 s) rather than only at times the rat pressed a lever. These models produced poor fits, because of fluctuations in overall rate of responding that appeared to be independent of the reward sequence.

The models described next differed in how Equation 1 was evaluated, corresponding to different generative models for the temporal dynamics of reward probability.

### Diffusion Model

The diffusion model assumes a generative model in which the reward probability for each lever evolves according to a diffusion process. More precisely, let 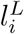 and 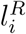 be the log-odds of the reward probabilities for the two levers at time *t*_*i*_, so that for each lever *X* (where *X* stands for *L* or *R*),

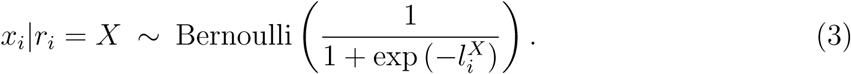

Then 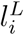 and 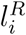 are taken to be governed by independent Wiener diffusion processes with diffusion rate *σ*^2^. Thus for each lever, the distribution of 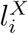 conditioned on 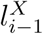 is a Gaussian with mean 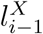 and variance proportional to the elapsed time, *t*_*i*_ *−t*_*i-*1_:

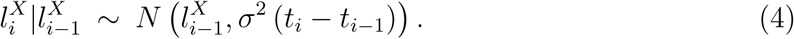

Representing the dynamics in terms of log-odds is convenient because it keeps the probability in the range [0, 1]. The model could equivalently be written as a non-homogenous diffusion process in the reward probability.

For both levers, the prior distribution on 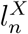 (i.e., before *x*_*n*_ is observed), is given by applying the diffusion dynamics to the posterior on 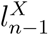:

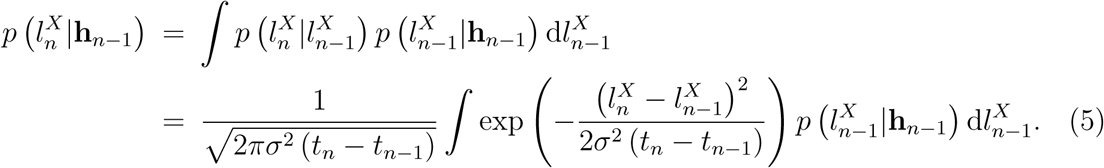

The expected reward value of each lever is obtained from this prior, as

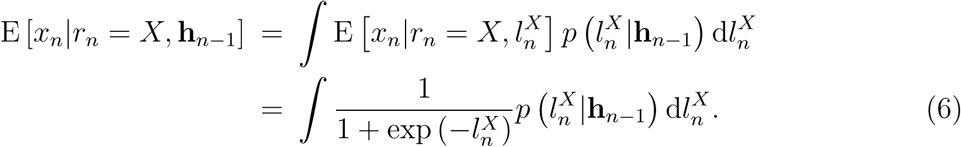

The posterior distributions on 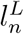 and 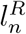 after *r*_*n*_ and *x*_*n*_ have been determined are obtained from Bayes’ rule. If *r*_*n*_ = *L*, then the posteriors are given by

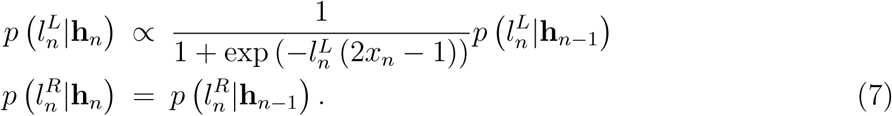

If *r*_*n*_ = *R* then Equation 7 is reversed, in that 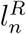 is updated and 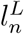 is not.

Iterating Equations 5 through 7 enables derivation of model predictions at all choice points. The free parameters of the diffusion model are the inverse temperature (*ϕ*), the response bias (*b*), and the diffusion rate (*σ*).

### Changepoint Model

The changepoint model assumes a generative model in which the reward probability for each lever evolves according to a changepoint process. Changepoints are assumed to be independent for the two levers, each generated by a Poisson process with rate parameter *λ*. Between changepoints for a lever, the reward probability for that lever is constant.

Denote the reward probabilities for the two levers at time *t*_*i*_ by 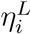 and 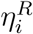, so that for each lever *X*,

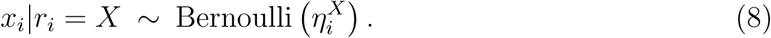

Let 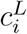 and 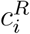 indicate whether one or more changepoints occur between times *t*_*i−*1_ and *t*_*i*_ (for the left and right levers, respectively). The Poisson assumption implies that for each lever,

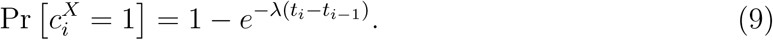

For each lever, the distribution of 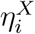 conditioned on 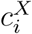 and 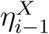 is given by

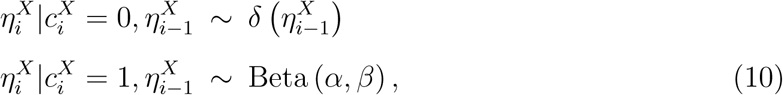

where *δ* is the Dirac delta function, and *α* and *β* are free repameters defining the distribution of *η* following a changepoint (taken to be a beta distribution).

The expected reward value of each lever *X* is equal to the mean of the prior distribution for *η*^*X*^:

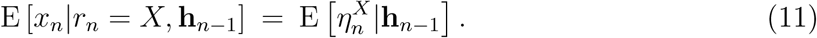

The prior distributions for *η*^*L*^ and *η*^*R*^ can be calculated by defining 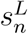 and 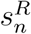 as the number of total lever presses since the last changepoint on each lever, prior to *t*_*n*_ (1):

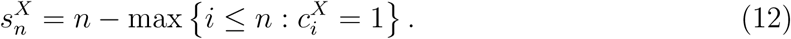

By convention we define 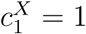, so that the maximal value of 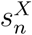 is *n −* 1.

Conditioned on 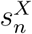, the prior distribution for 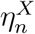 is given by Bayesian inference over the last 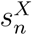 outcomes, with Beta (*α, β*) as the prior:

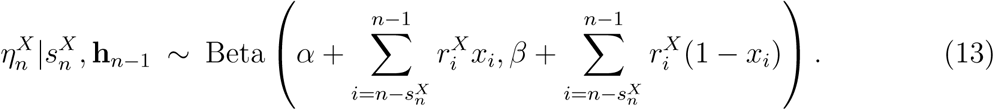

Here 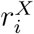 equals 1 if *r*_*i*_ = *X* and 0 otherwise. The sums represent the numbers of rewarded and nonrewarded presses of lever *X* out of the last 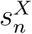 total lever presses (if 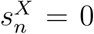 then these sums are taken to be empty).

The prior distribution for 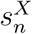 is obtained from the posterior on 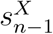 by shifting the values by 1 and applying the probability of an intervening changepoint:

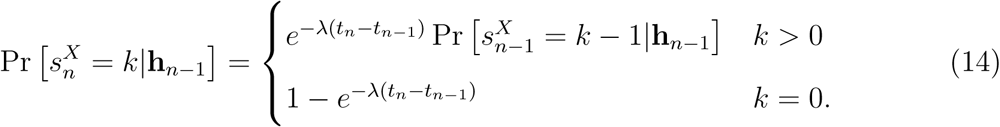

The posterior distribution for 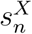 can be obtained from Bayes’ rule. If *r*_*n*_ = *X*, then the posterior for 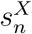 is given by

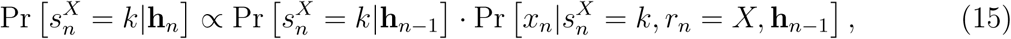

where the likelihood is obtained using Equation 13:

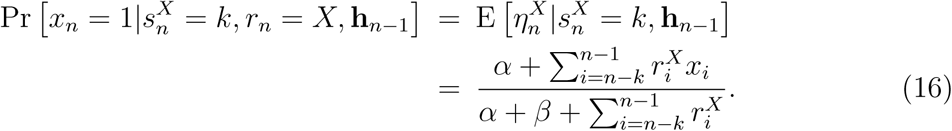

The posterior for the unpressed lever is identical to its prior. That is, if *r*_*n*_ ≠ *X*, then

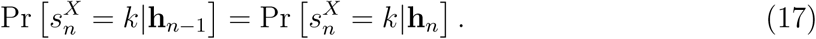

Iterating Equations 14, 15, and 17 enables calculation of the prior distributions for 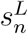 and 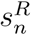 at all choice points. The expected reward value of each lever *X* can be obtained by summing this prior for 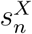:

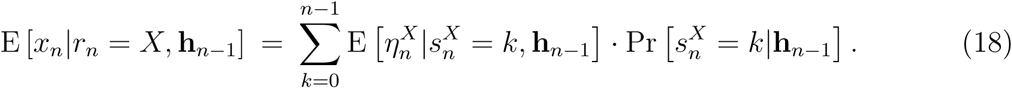

The free parameters of the changepoint model are the inverse temperature (*ϕ*), the response bias (*b*), the changepoint rate (*λ*), and the parameters of the reward-probability distribution following a changepoint (*α* and *β*).

### Full Model

The full model evaluates Equation 1 by combining the predictions of the diffusion and changepoint models. The combination is Bayesian except for a recency bias in evaluating the likelihood of each subsystem. Specifically, the subsystems’ predictions are weighted according to

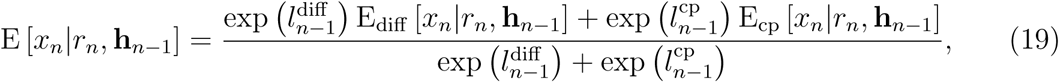

where E_diff_ and E_cp_ are the expectations of the two subsystems, taken from Equations 6 and 18.

The mixture weights are obtained from a running average of each subsystem’s log-likelihood of the results of past lever presses:

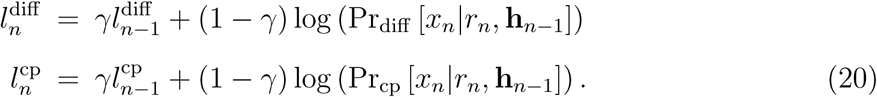

The *γ* parameter controls the rate of decay of the influence of past events. When *γ* = 1, 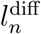 and 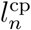 are the exact log-likelihoods of the rat’s full experience in the task, and the model becomes an exact hierarchical Bayesian model. When *γ <* 1, the model assumes that recent performance is more important in determining how much each subsystem should drive behavior. Equation 20 produces exponential decay, which approximately matches the optimal decay profile under both diffusion and changepoint dynamics (2). Thus the hierarchical model is agnostic about the form of nonstationarity in the relative performance of the two subsystems.

The free parameters of the hierarchical model are the parameters of the diffusion and changepoint models (*ϕ, b, σ, λ, α, β*) and the decay rate (*γ*).

### Model Evaluation

Models were simulated on each rat’s data from phases 1 and 2 combined. Three different models were fit to the individual data of each subject: the full model, a model in which only the diffusion model contributed to action selection in phase 2, and a model in which only the changepoint model contributed to action selection in phase 2. In phase 1, all three models derived reward expectations from Equation 19. In phase 2, the full model continued to use Equation 19, the diffusion model used Equation 6, and the changepoint model used Equation 18.

We evaluated the fit of each model by calculating its log-likelihood, that is, the sum of the log of the probabilities the model assigned to each lever press from Equation 2. Models were evaluated only in their predictions for which lever the rat pressed, conditioned on when each press occurred. That is, we attempted to fit the time course of each rat’s relative preference between the two levers, setting aside variation in overall response rate. Video data indicated that distractions and other activities (e.g., grooming) modulated the rats’ engagement in the task in a manner the models should not be expected to capture. Therefore, at each time the rat pressed a lever, the model was queried for its current response probability for pressing left versus right.

For simulating the diffusion model, the ranges of *l*^*L*^ and *l*^*R*^ were discretized into 401 bins each. The iterative prior and posterior distributions were calculated on this discretized representation. Simulation of the changepoint model was based on maintaining the complete distributions of *s*^*L*^ and *s*^*R*^.

## References

1. Chater, N., & Manning, C. D. (2006). Probabilistic models of language processing and acquisition. Trends in Cognitive Sciences, 10, 7, 335–44.

2. Xu, F., & Tenenbaum, J. B. (2007). Word learning as Bayesian inference. Psychological Review, 114, 245–272.

3. Anderson, J. (1991). The adaptive nature of human categorization. Psychological Review, 98, 409–429.

4. Griffiths, T. L., & Tenenbaum, J. B. (2009). Theory-based causal induction. Psychological Review, 116, 661–716.

5. Chater, N., & Oaksford, M. (1999). The probability heuristics model of syllogistic reasoning. Cognitive Psychology, 38, 191–258.

6. Goodman, N. (1955). Fact, Fiction, and Forecast. Third chapter: The new riddle of induction. Cambridge, MA: Harvard University Press.

7. Hume, D. (1748/1975). An Enquiry concerning Human Understanding. In Enquiries concerning Human Understanding and concerning the Principles of Morals, ed. L. A. Selby-Bigge and P. H. Nidditch, 5–165. Oxford: Clarendon.

8. Wolpert, D. (1996). The lack of a priori distinctions between learning algorithms. Neural Computation, 1341–1390.

9. Wolpert, D.H., Macready, W.G. (1997). No Free Lunch Theorems for Optimization. IEEE Trans Evol Comput 1, 1, 67–82.

10. Bowers, J.S., & Davis, C.J. (2012). Bayesian just-so stories in psychology and neuroscience. Psychological Bulletin, 138, 389–414.

11. Jones M., & Love B. (2011). Bayesian fundamentalism or enlightenment? On the explanatory status and theoretical contributions of Bayesian models of cognition. Behavioral and Brain Sciences 34, 169–188.

12. Savage, L.J. (1954). The foundations of statistics. John Wiley/Dover.

13. Gigerenzer, G., & Brighton, H. (2009). Homo heuristicus: Why biased minds make better inferences. Topics in Cognitive Science, 1, 107–143.

14. Kemp, C., & Tenenbaum, J. B. (2008). The discovery of structural form. Proceedings of the National Academy of Sciences of the United States of America, 105, 10687–10692.

15. Tenenbaum, J.B., Kemp, C., Griffiths, T.L., & Goodman, N.D. (2011). How to Grow a Mind: Statistics, Structure, and Abstraction. Science, 331(6022), 1279–1285.

16. Ashby F.G., Alfonso-Reese, L.A., Turken A.U., & Waldron E.M. (1998). A neuropsychological theory of multiple systems in category learning. Psychological Review, 105, 442–481.

17. Jacobs, R.A., Jordan, M.I., Nowlan, S.J., & Hinton, G.E. (1991). Adaptive mixtures of local experts. Neural Computation, 3, 79–87.

18. Merton, R.C. (1998). Applications of option-pricing theory: Twenty-five years later. American Economic Review, 88, 323–349.

19. Estes, W.K. (1950). Toward a statistical theory of learning. Psychological Review, 57, 94–107.

20. Rescorla, R.A., & Wagner, A.R. (1972). A theory of Pavlovian conditioning: Variations in the effectiveness of reinforcement and nonreinforcement. In Classical Conditioning II (ed. A.H. Black, W.F. Prokasy), 64–88. New York: Appleton-Century-Crofts.

21. Sutton, R.S., & Barto, A.G. (1998). Reinforcement Learning. Cambridge, MA: MIT Press.

22. Wilder, M., Jones, M., & Mozer, M.C. (2010). Sequential effects reflect parallel learning of multiple environmental regularities. In Y. Bengio, D. Schuurmans, J. Lafferty, C. K. I. Williams, & A. Culotta (Eds.), Advances in Neural Information Processing Systems, 23, 2053–2061.

23. Yu, A. & Cohen, J. (2009). Sequential effects: Superstition or rational behavior? In Y. Bengio, D. Schuurmans, J. Lafferty, C. K. I. Williams, & A. Culotta (Eds.), Advances in Neural Information Processing Systems, 22, 1873–1880.

24. Brown, S. D., & Steyvers, M. (2009). Detecting and predicting changes. Cognitive Psychology, 58, 49–67.

25. Gallistel, C.R., Mark, T.A., King, A.P., & Latham, P.E. (2001). The rat approximates an ideal detector of changes in rates of reward: implications for the law of effect. Journal of experimental psychology: Animal behavior processes, 27, 354–372.

26. Herrnstein, R. J., Rachlin, H., & Laibson, D. I. (2000). The Matching Law. Cambridge, MA: Harvard University Press.

27. Pauli, W. M., Clark, A. D., Guenther, H., O’Reilly, R. C., & Rudy, J. W. (2012). Inhibiting pkm_zeta reveals dorsal lateral and dorsal medial striatum store the different memories needed to support adaptive behavior. Learning and Memory, 19, 307–314.

28. Yin, H.H., Knowlton, B.J., & Balleine, B.W. (2005a). Blockade of NMDA receptors in the dorsomedial striatum prevents action-outcome learning in instrumental conditioning. European Journal of Neuroscience, 22, 505–512.

29. Yin, H.H., Knowlton, B.J., & Balleine, B.W. (2005b). The role of the dorsomedial striatum in instrumental conditioning. European Journal Neuroscience, 22, 513–523.

30. Yin, H.H., Knowlton, B.J., & Balleine, B.W. (2004). Lesions of dorsolateral striatum preserve outcome expectancy but disrupt habit formation in instrumental learning. European Journal of Neuroscience, 19, 181–189.

31. Hazy, T. E., Frank, M. J. & O’Reilly, R. C. (2007). Towards an executive without a homunculus: computational models of the prefrontal cortex/basal ganglia system. Philosophical Transactions of the Royal Society B, 362, 1601–1613.

32. Sacktor, T.C. (2011). How does PKMzeta maintain long-term memory? Nature Reviews Neuroscience, 12, 9–15.

33. Heyman G.M. (1979). A Markov model description of changeover probabilities on concurrent variable-interval schedules. Journal of the Experimental Analysis of Behavior, 31, 41–51.

34. Marr, D. (1982). Vision: A Computational Investigation into the Human Representation and Processing of Visual Information. New York: Freeman

35. Ashby, F. G., & Maddox, W. T. (2005). Human category learning. Annual Review of Psychology, 56, 149–178.

36. Daw, N.D., Niv, Y., & Dayan, P. (2005). Uncertainty-based competition between prefrontal and dorsolateral striatal systems for behavioral control. Nature Neuroscience, 8, 1704–1711.

37. Bellman, R. E. (1957). Dynamic Programming. Princeton, NJ: Princeton University Press.

38. Watkins, C. J. C. H., & Dayan, P. (1992). Q-Learning. Machine Learning, 8, 279–292.

39. Rougier, N. P., Noelle, D., Braver, T. S., Cohen, J. D., & O’Reilly, R. C. (2005). Prefrontal cortex and the flexibility of cognitive control: Rules without symbols. Proceedings of the National Academy of Sciences, 102, 7338–7343.

40. Featherstone, R.E., & McDonald, R.J. (2004). Dorsal striatum and stimulus-response learning: Lesions of the dorsolateral, but not dorsomedial, striatum impair acquisition of a stimulus-response-based instrumental discrimination task, while sparing conditioned place preference learning. Neuroscience, 124, 23–31.

## References

1. Adams, R. P., & MacKay, D. J. (2007). Bayesian online changepoint detection. Technical report, University of Cambridge, Cambridge, UK.

2. Yu, A. & Cohen, J. (2009). Sequential effects: Superstition or rational behavior? In Y. Bengio, D. Schuurmans, J. Lafferty, C. K. I. Williams, & A. Culotta (Eds.), Advances in Neural Information Processing Systems, 22, 1873–1880.

